# ROS Mediated the Photodynamic Antibacterial of *Vibrio vulnificus* by a DNA-Targeted Molecular Rotor Probe

**DOI:** 10.1101/2024.04.03.587862

**Authors:** Haiyun Yue, Qing Bian, Xintong Li, Chao Yu, Chao Chen, Kangnan Wang, Yuan Cao

**Affiliations:** Department of Basic Medical Sciences, The 960th Hospital of PLA, Jinan 250031, China; State Key Laboratory of Crystal Materials, Shandong University, Jinan 250100, China

**Keywords:** *Vibrio vulnificus*, Antibacterial, Photosensitizer, Photodynamic

## Abstract

*Vibrio vulnificus*, a highly pathogenic Gram-negative bacterium, is capable of inducing sepsis, necrotizing fasciitis, and skin and soft tissue infections through contact with wounds. Currently, the majority of *V. vulnificus* strains have developed resistance to multiple drugs, highlighting the critical necessity for the development of novel therapeutics capable of effectively targeting and eradicating this bacterium. In recent years, material molecules have emerged as promising antimicrobial agents. This study introduces a novel molecular fluorescent probe, BDTP, which demonstrates a wide-ranging antimicrobial effect against both Gram-negative and Gram-positive bacteria while exhibiting minimal toxicity to normal mammalian cells. Of particular significance is BDTP’s ability to rapidly detect V. vulnificus, bind to bacterial DNA, and exhibit fluorescence monitoring behavior. Furthermore, BDTP displays enhanced photodynamic antibacterial activity when exposed to white light irradiation. Under a low dose of white light (15mW cm^-2^), the killing efficiency of *V. vulnificus* irradiated with 4uM for 10 minutes was more than 99.8%. Moreover, it could significantly inhibit and eliminate the biofilm formed by *V. vulnificus*. It can induce the production of reactive oxygen species (ROS) in *V. vulnificus* cells, leading to bacterial cell damage. More importantly, BDTP significantly promoted the healing of infected wounds in an animal model of *V. vulnificus* infection. Therefore, BDTP shows great promise as a potent antibacterial agent against *V. vulnificus* infection.

## 1. Introduction

*Vibrio vulnificus* (*V. vulnificus*) is a kind of halophilic, gram-negative bacterium vibrio, belongs to the conditional pathogenic bacteria. It was widely existed in coastal waters worldwide such as the China, USA, Japan, South Korea, and Mexico^[1]^. Most commonly, humans get the infection from ingesting contaminated seafood or from wounds contact with seawater^[2]^. The main clinical symptoms were severe gastroenteritis, wound infection, cellulitis and highly fatal septicemia^[3]^. In particular, individuals suffering from liver disease, cancer and immunocompromised individuals are more likely to develop fatal wound infection and primary sepsis once they are infected with *V. vulnificus*^[4]^. Patients with *V. vulnificus* infection tend to have a rapidly progressive clinical course, with more than 50% of patients dying within 48 hours of hospitalization^[5]^. With global warming, the distribution of Vibrio vulnificus has further expanded, causing an increase in the number of infections year by year^[4, 6]^. However, the misuse of antibiotics in the past few decades has led to an increase in *V. vulnificus* resistance to antibiotics worldwide and significantly reduced treatment efficacy^[7, 8]^. Therefore, there is an urgent need to develop drugs without causing antimicrobial resistance but can quickly recognize and kill *V. vulnificus*.

Antimicrobial photodynamic therapy (aPDT) has become a promising method in antibacterial applications due to its non-invasive treatment, simple operation, low cost and low resistance^[9-13]^. In the process of aPDT, light excited the photosensitizers (PSs) to generate a large amount of reactive oxygen species (ROS) which can cause oxidative damage to the cell membrane and DNA of bacteria, and eventually induce the death of bacteria^[14, 15]^. In addition, PDT can also lead to the ablation of bacterial biofilms^[15, 16]^. It is well known that the formation of biofilm can counter the effects of drugs and the host immune system, increasing the ability of bacteria to infect the human body^[17]^. At the same time, it also makes the bacteria encapsulated in biofilm resistant to antibiotics up to 10 to 1000 times, which greatly increases the difficulty of antibacterial therapy^[18]^. Therefore, the inhibition or elimination of bacterial biofilm is an important part of the antibacterial process. In conclusion, PDT is considered as an attractive antibacterial strategy.

In recent years, material molecules have shown great potential as antimicrobial agents^[19, 20]^. However, because of the structural differences between Gram-positive and Gram-negative bacteria, most photosensitizer (PS) molecules only exhibit antimicrobial activity against Gram-positive bacteria but failed to affect Gram-positive bacteria^[13, 21]^. Hence, this largely limits the practical application of photosensitizers. Here, we present a molecular fluorescent probe (bacteria DNA-Targeted Photosensitizer, BDTP) with positively charged pyridine groups as the two arms, and the hydrophilic heads not only improve the molecular rotor effect, but also help the probe to bind to negatively charged bio-components (e.g., phospholipid and DNA). BDTP can rapidly recognize both Gram-positive and Gram-negative bacteria and binds to the DNA of the bacteria with broad spectrum antimicrobial activity and fluorescence monitoring behavior. Under white light irradiation, BDTP converts light energy into ROS, which leads to oxidative DNA damage and further bacterial rupture. In this study, we extensively investigated the targeting and phototherapeutic effects of BDTP on *V. vulnificus* and biofilm through in vivo and in vitro experiments.

## 2. Results and discussion

### 2.1 Structure and photophysical properties

In this study, we screened and prepared a photosensitizer named BDTP by characterizing the features of bacterial and eukaryotic cell structures^[22]^. The photosensitizer is equipped with two positively charged pyridinium salt units, which enhance the binding ability of the photosensitizer to DNA and improve the electron-absorbing properties of the molecule, resulting in a redder fluorescence emission. Additionally, a relatively lipophilic cinnamoyl unit was introduced into the symmetric segment of the photosensitizer to ensure prolonged retention on the membrane structure of eukaryotic cells. As shown in Figure 1A, the photosensitizer exhibits maximum absorption and fluorescence emission peaks around 420 nm and 651 nm, respectively. Upon addition of approximately 40 equivalents of double-stranded DNA in Tris-HCl solution, the fluorescence intensity of the photosensitizer increased by about 150-fold (Figure 1B), indicating an extremely sensitive fluorescence response of the photosensitizer to DNA. Furthermore, we conducted dynamic simulations of the binding mode between the photosensitizer and DNA (PDB: 5SWD) using a semi-flexible docking approach^[23]^. Experimental results showed in figure 1C-D demonstrated that the photosensitizer can be embedded into the major groove of DNA through strong electrostatic interactions while showing weaker interactions with the proteins surrounding DNA.

**Figure 1.**
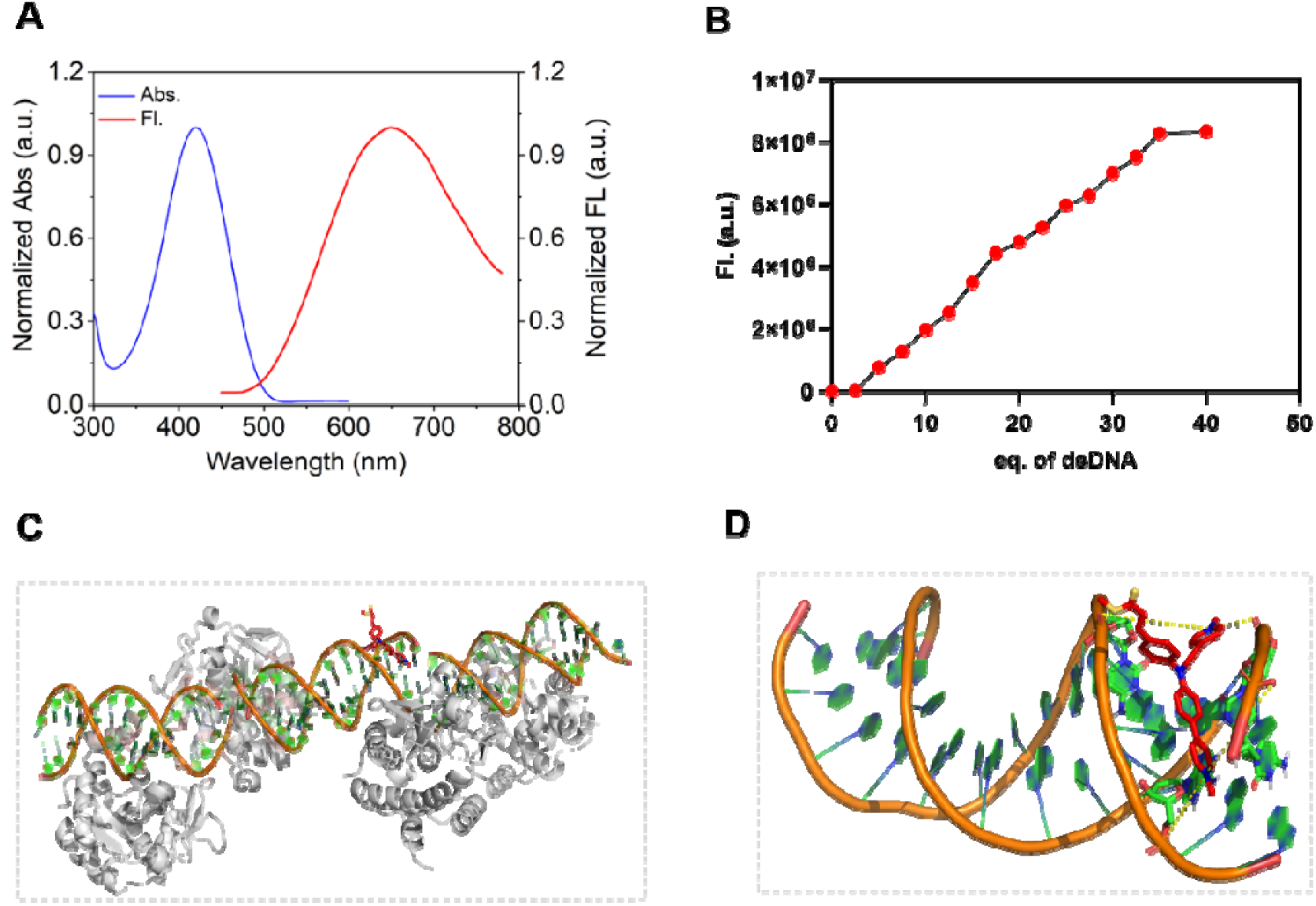

Next, we conducted a preliminary evaluation of BDTP as a type I/II photosensitizer in terms of generating reactive oxygen species (ROS). Initially, we utilized ABDA as a probe to assess the potential of BDTP to generate singlet oxygen (^1O_2) through the type II pathway. ABDA undergoes oxidation in the presence of ^1O_2, leading to a decrease in absorption at 378 nm. When exposed to white light, BDTP induced the degradation of ABDA by 55.78 %, outperforming the commercial photosensitizer Rose Bengal (RB) (Figure 2A). We further employed dihydrorhodamine 123 (DHR123) as a selective fluorescence probe for superoxide anion (O_2•^–). Upon reacting with O_2•^–, DHR123 emits strong green fluorescence at 530 nm when excited at 500 nm. The control group containing only DHR123 exhibited slight fluctuations at 530 nm after 300 seconds of irradiation, indicating that white light alone does not oxidize DHR123 (Figure 2B). However, when DHR123 and BDTP were irradiated in PBS for 300 seconds, the fluorescence of DHR123 increased by 20-fold (Figure 2B). Under the same conditions, the fluorescence of BDTP-induced DCF-DA increased by 10-fold (Figure 2C). Additionally, we utilized hydroxyphenyl fluorescein (HPF) to evaluate the effect of BDTP on generating hydroxyl radicals, as shown in Figure 2D. After 10 minutes of white light irradiation, a significant enhancement in fluorescence signal attributed to BDTP was observed. Overall, BDTP demonstrated the ability to generate type I and type II ROS. This lays the foundation for the photosensitizer’s potential to damage bacterial nucleic acids effectively.

**Figure 2.**
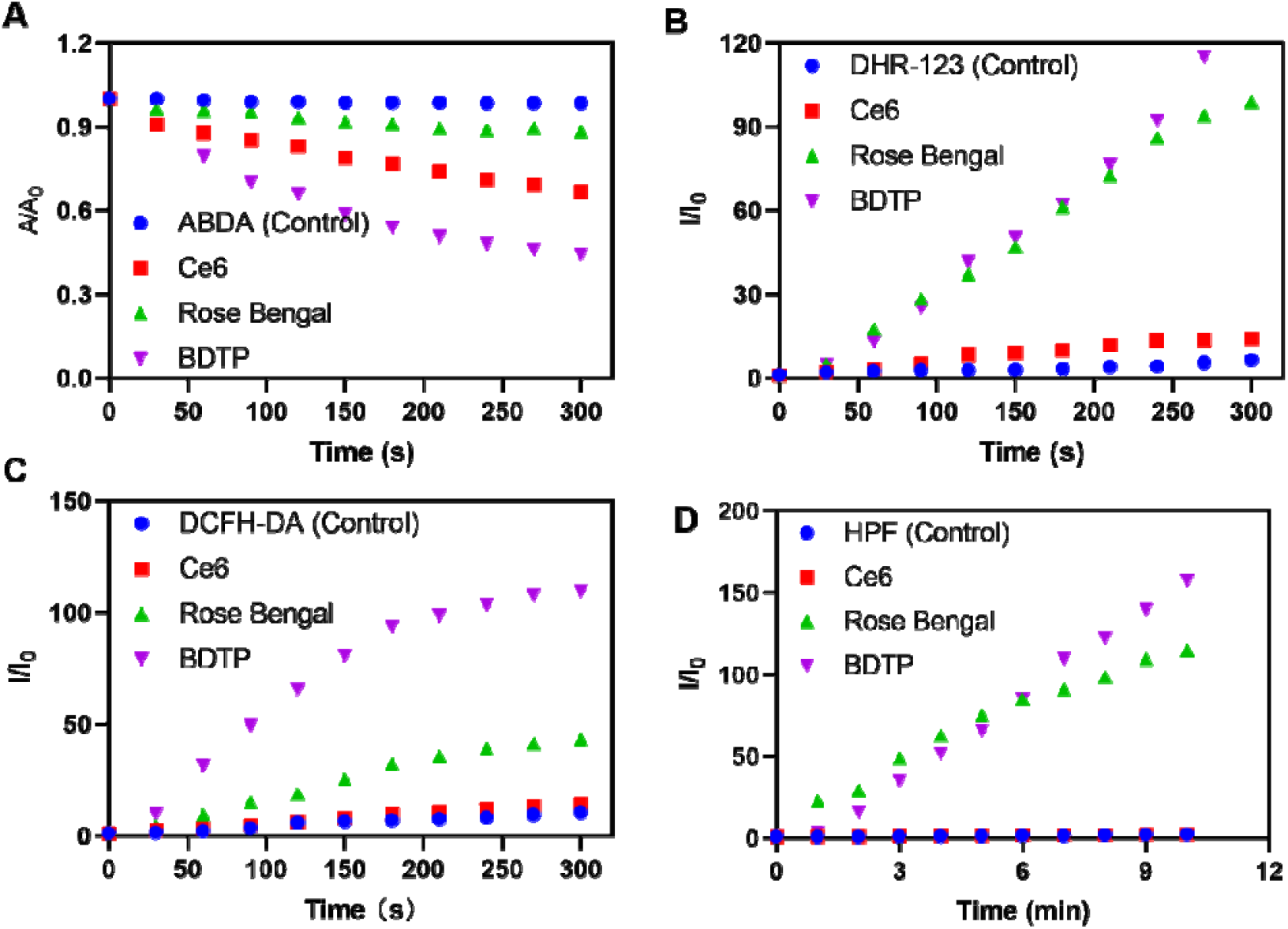

### 2.2 Antibacterial Activity of BDTP Against Free Bacteria

The antimicrobial activity of BDTP against the Gram-positive bacteria *Staphylococcus aureus* and the Gram-negative bacteria *Escherichia coli* was assessed by measuring the relative number of colony-forming units (CFUs) on AGAR plates. As shown in Figure3A, significant antibacterial ability of BDTP with dark treatment could be observed compared with the control group (0μM BDTP). In the BDTP+ dark treatment group, the relative amount of CFUs gradually decreased with the increase of BDTP concentration. As shown in Figure 3B, the relative viability of *E. coli* decreased to 0% under treatment with BDTP at the concentration of 64μM without light irradiation, while the relative viability of *S. aureus* decreased to 0% at the concentration of 16μM (Figure 3C). BDTP with white light irradiation (15mW cm^-2^, 30 min) treatment showed more obvious antibacterial property. The relative viability of *E. coli* decreased to 0% at 16μM of BDTP with white light irradiation (15mW cm^-2^, 30 min). Furthermore, the relative amount of CFUs of *S. aureus* decreased to 0% at 4μM of BDTP with the same light treatment (Figure 3B, C). However, the light treatment without BDTP,*E. coli* and *S. aureus* did not show any significant change contract with the dark treatment. Which indicated that light irradiation itself does not lead to cell ablation.

**Figure 3.**
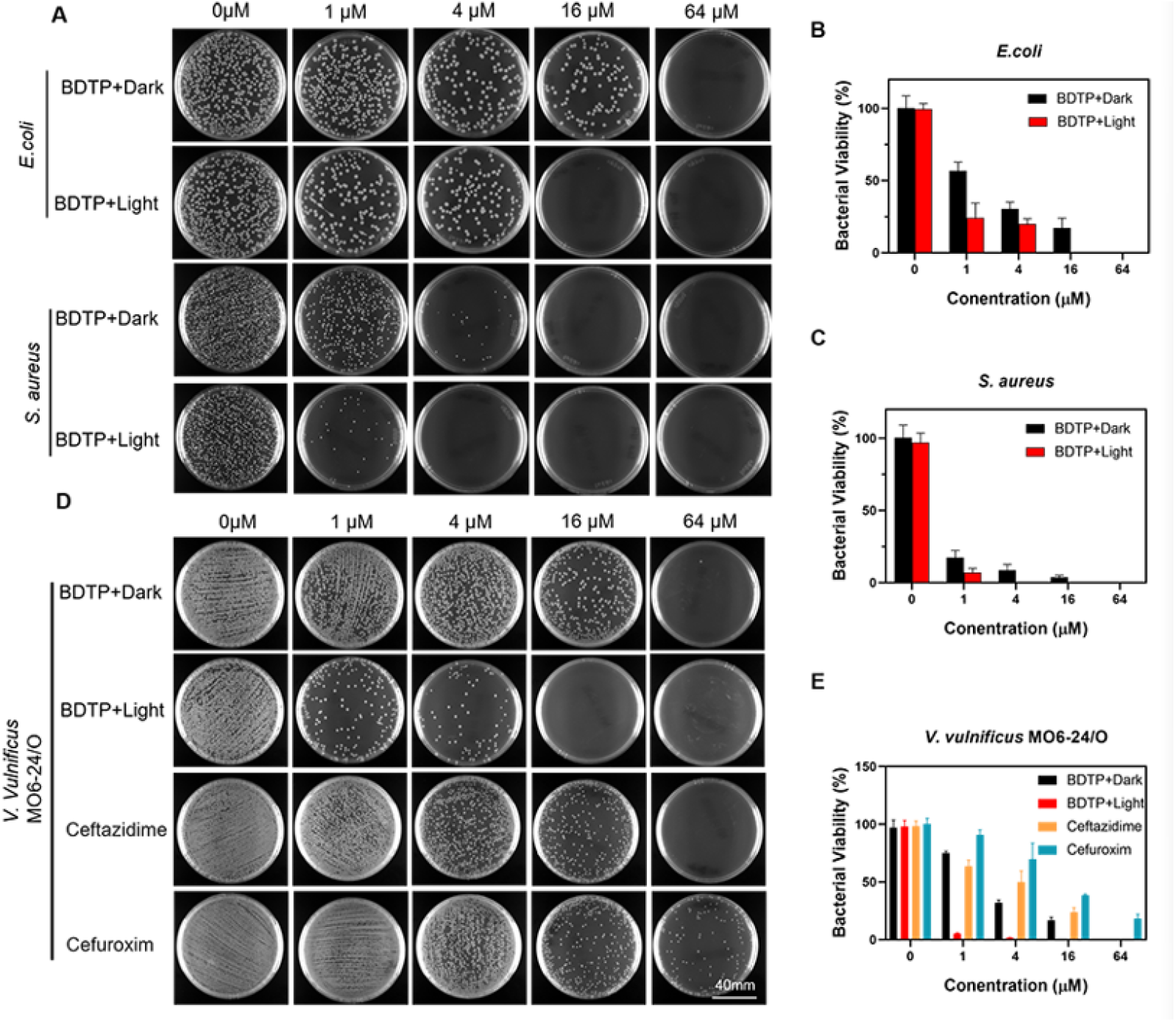
Evaluation of the antibacterial activity of BDTP. (A) Antibacterial activity study by different concentration of BDTP against gram-negative bacteria *E. coli* and the gram-positive bacteria *S. aureus* with or without light irradiation (15mW cm^−2^, 30min). (B-C) Inhibition of bacterial viability of BDTP under different concentrations with or without light irradiation. (n=3, mean±SD) (D) Antibacterial activity study by different concentration of BDTP against *V. vulnificus* with or without light irradiation (15mW cm^−2^, 30min). (E) Inhibition of bacterial viability of *V. vulnificus* under different concentrations with or without light irradiation. (n=3, mean±SD)

Since BDTP showed excellent inhibition against *E. coli* and *S. aureus*, and because of the high pathogenicity and high mortality of *V. vulnificus*, as well as its extensive of drug resistance^[24]^, the antibacterial activity of BDTP against *V. vulnificus* was investigated. The relative amount of CFUs of *V. vulnificus* was measured. Figure 3D, E showed that after BDTP treatment with dark, the amount of CFUs of *V. vulnificus* decreased with the increase of BDTP concentration, and under the treatment of BDTP with the concentration of 64μM, the bacterial viability was completed disappeared. Which was similar to the antibacterial effect of ceftazidime and cefuroxime on *V. vulnificus*. However, in the BDTP with light treatment group, the relative bacterial viability of *V. vulnificus* was significantly lower than that in the dark treatment group, showing a better PDT antibacterial effect. As shown in Figure 3E, the relative bacterial viability of *V. vulnificus* after treatment of 1μM BDTP wiht light was decreased to 5%, and after 4μM of BDTP with light treatment, the relative viability of *V. vulnificus* decreased to 1.5%.

### 2.3 Biofilm Inhibition and Eradication Activity of BDTP Toward *V. vulnificus*

It was found that the pathogenicity of *V. vulnificus* was correlate with biofilm formation ability^[25]^. Bacterial biofilms with dense physical structure and polysaccharide-and nucleic acid-rich extracellular polymeric substance, therefore, they have the ability to resistant to antibiotic permeability and protect them from oxygen free radicals, disinfectants, pH changes, low nutrients^[26]^. In addition, the formation of biofilms promotes the rapid colonization and invasion of pathogens in the host and facilitates evasion of host defense mechanisms^[27]^. Based on the study of the antibacterial activity of BDTP against free bacteria, we investigated the inhibitory and eliminative effects of BDTP on *V. vulnificus* biofilms.

*V. vulnificus* biofilms were prepared by inoculations of bacterial solution containing different concentrations of BDTP into 96-well plates. The inhibition of BDTP was evaluated by crystal violet staining. The crystal violet staining results shows that BDTP has obvious anti-biofilm effect, what’s more interesting is that the anti-biofilm efficiency of BDTP with white light irradiation is much higher than PDT or antibiotic treatment alone (Figure 4A, B). Moreover, BDTP at the concentration of 16μM with white light irradiation (15mW cm^-2^, 30 min) suppressed the biofilm formation up to 95.85% (Figure 4C), which demonstrated the effectiveness of PDT activated of BDTP. In addition, CLSM results showed a significant reduction of green fluorescence of SYTO9 labeled the bacteria in the *V. vulnificus* biofilm after treatment with BDTP and light irradiation (Figure 4D, E), which suggested that the biofilm formation was suppressed remarkable under the PDT treatment.

**Figure 4.**
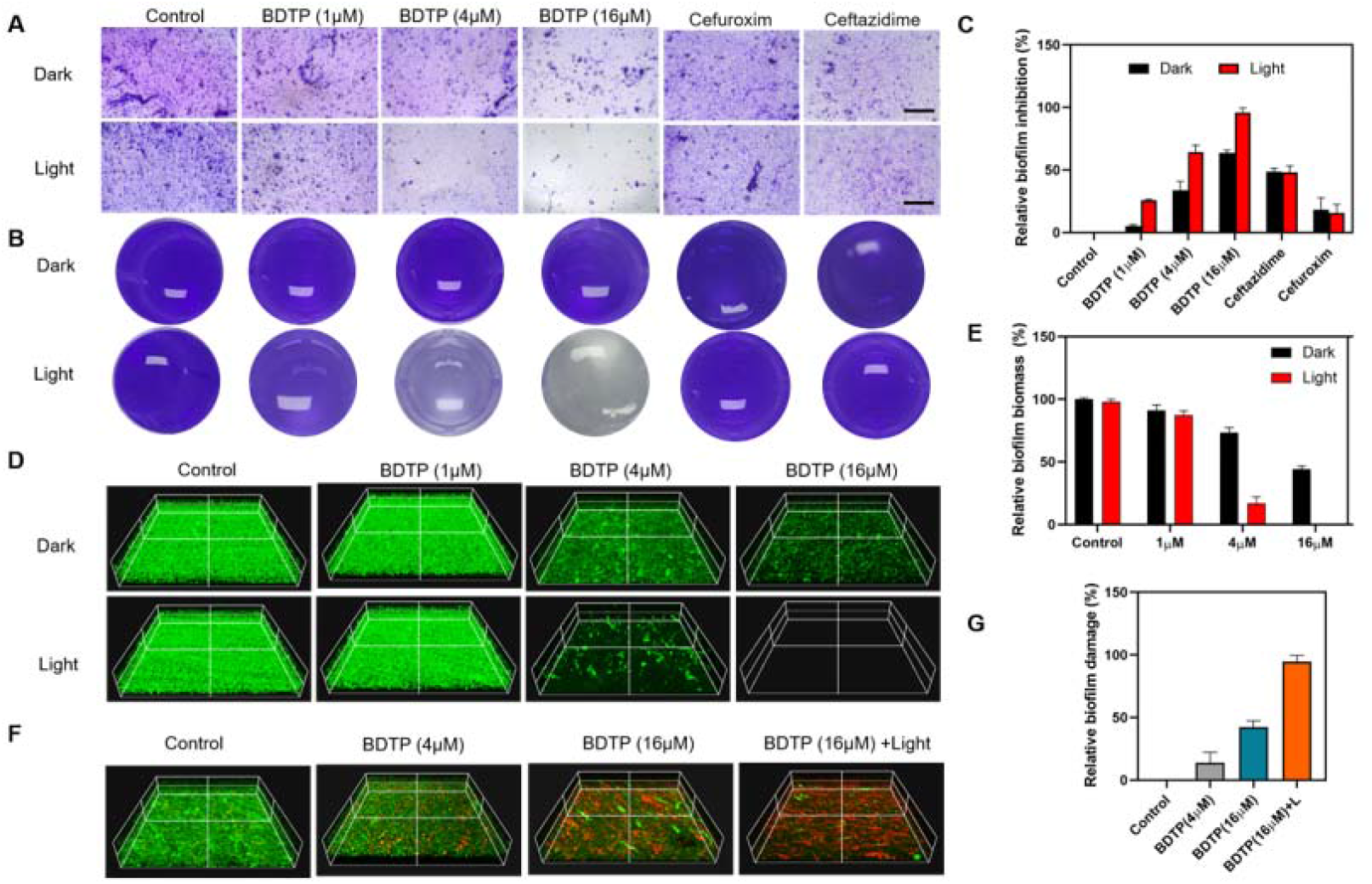
Antibiofilm properties of BDTP. (A-B) The photographs of *V. vulnificus* biofilm stained by crystal violet after different treatments. Scale bar is 200μm. (C) Relative biofilm inhibition ratio of BDTP for *V. vulnificus* after various treatments. (n=3, mean±SD) (D) Representative CLSM images of 3D reconstructions of *V. vulnificus* biofilm stained by SYTO9. (E) Relative fluorescence quantity of the CLSM results. (n=3, mean±SD) (F) 3D reconstructions CLSM images of *V. vulnificus* biofilm stained by SYTO9 and PI. (G) Relative biofilm damage ratio of BDTP for *V. vulnificus* after various treatments. (n=3, mean±SD)

For explore the eradication of BDTP toward *V. vulnificus* biofilm, we then treated the biofilm with BDTP for 30min, then give the dark or light irradiation (15mW cm^-2^, 30min), the CLSM images demonstrated the ability of BDTP to eradicate biofilms. As showed in Figure 4F, after the treatment of BDTP without light irradiation, the viability of the biofilm was decreased dramatically with the increase of BDTP concentration. Moreover, after the treatment with BDTP and light irradiation, the red fluorescence intensity of PI label the dead bacterial was significantly enhanced. This indicated that the photodynamic activity of BDTP causes great damage to the *V. vulnificus* biofilm. These results suggested that BDTP has good anti-biofilm formation ability and eradication activity. Therefore, it is expected to be a candidate molecule for the photodynamic treatment of anti-biofilm of *V. vulnificus*.

### 2.4 BDTP can identify the bacteria and colocalize with the DNA

Since BDTP shows good antibacterial and anti-biofilm function, we investigated the localization of BDTP in bacteria. *E. coli* and *V. vulnificus* as gram-negative bacterial models and *S. aureus* as gram-positive bacterial model incubated with 4μM BDTP for 10min respectively, and then incubated with Hoechst 33342 to stain the bacterial DNA. The CLSM images showed that Hoechst 33342 stained bacterial DNA with green fluorescence, and the red fluorescence was emission from the bacterial stained by BDTP. The results clearly showed that the BDTP red fluorescence coincided perfectly with the Hoechst 33342 green fluorescence (Figure 5A), suggesting that BDTP could effectively anchor to the bacterial DNA. In the past few decades, many organic fluorescent probes were used for bacterial imaging ^[28]^. The fluorescent labeling of bacterial DNA also provides a direction for studying the mechanism of bacterial resistance^[29]^.

**Figure 5.**
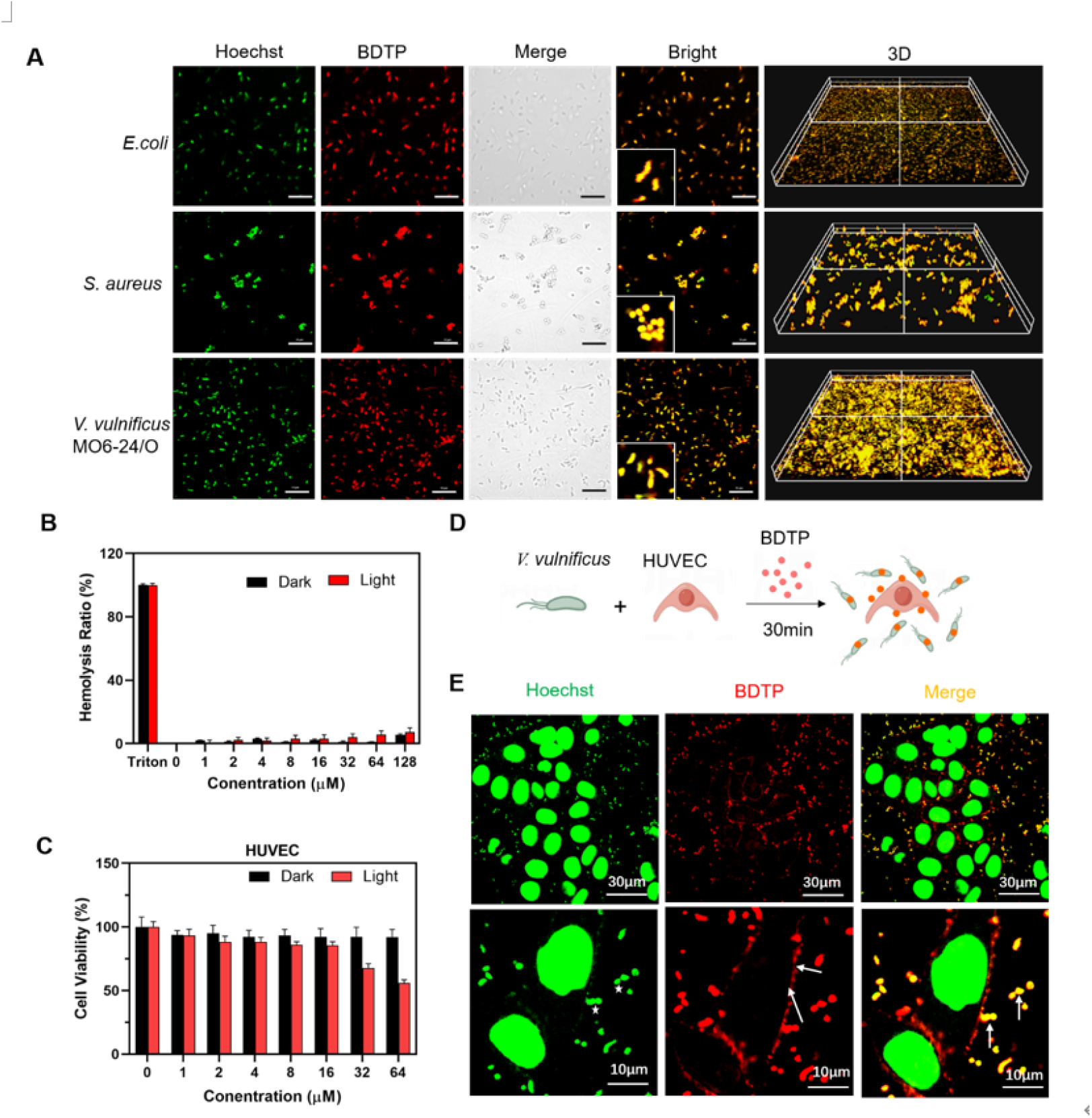
Detection of BDTP localization in bacteria and cells. (A) CLSM images of bacteria treated with BDTP. (B) Cell viability of HUVEC cells treated with different concentrations of BDTP, with or without irradiation. (n=3, mean±SD) (C) Hemolysis ratio of RBCs treated with different concentrations of BDTP, with or without irradiation. (n=3, mean±SD) (D) Scheme of the BDTP labeled the bacterial and cells. (E) Representative fluorescence images of the co-culture treated with BDTP, green fluorescence represents DNA stained with Hoechst and red fluorescence represents BDTP. The images are representative of three independent experiments.

BDTP not only fluorescently labeled live bacteria with high specificity, but also showed low cytotoxicity and good hemocompatibility. As shown in Figure 5B, even when the concentration of BDTP was as high as 128 μM, the hemolysis ratios of BDTP was still less than 5% in the presence or absence of light, indicating that BDTP had good blood cell compatibility. In addition, the cytotoxicity of BDTP was investigated in the Human Umbilical Vein Endothelial Cells (HUVEC) via the cell counting kit-8 (CCK-8) assay. The results showed that the viability of cells was remains as high as 80% after the treatment of different concentration of BDTP(0-64μM) for 24 h in the dark. Even under the concentration as high as 64μM of BDTP treatment with white light irradiation (15mW cm^-2^, 30min), the relative viability of cells was still above 50% and did not cause significant cytotoxicity (>IC50, the minimal concentration causing 50% death of cells) (Figure 5C), indicating that BDTP has low cytotoxicity. Some biological imaging fluorescent probes, such as fluorescein derivatives, BODIPY derivatives, and SYTO-9TM, cannot be widely used due to the easy fluorescence quenching, high fluorescence background, and intense toxicity^[28, 30]^ºThe efficient bacterial DNA labeling activity and excellent biocompatibility of BDTP may be attributed to the water solubility and AIE activity properties of BDTP. BDTP maintains the molecular state in buffer, is radiation-free, and produces an ultra-low background signal in bioimaging even without washing.

Then the co-localization of BDTP with bacteria and cells was investigated to further explore the toxicity of BDTP. The scheme of the treatment for bacterial and cells was shown in Figure 5D. After co-culturing *V. vulnificus* and HUVEC cells for 30min, BDTP (4μM) was added to the co-culturing dish and incubated for 10min, and then incubated with Hoechst 33342 to stain the DNA. CLSM images showed that red fluorescence of BDTP rapidly bound to the bacterial cells and co-localized with the green fluorescence of *V. vulnificus* DNA. However, the HUVEC cells showed weak red fluorescence stained on the cell membrane but did not enter the cells (Figure 5E). Which probably contributed to the lower toxicity of BDTP.

### 2.5 Antibacterial Mechanism

In aPDT, photosensitizer molecules absorb the energy of photons and generate reactive oxygen species (ROS), which in turn cause oxidative damage to different bacterial structures and eventually kill bacterial pathogens^[31]^ºThe H_2_DCF-DA assay was used to determine the effect of BDTP on intracellular ROS production in *V. vulnificus* cells to further explore the antibacterial mechanism. Figure 6A-B showed that after treatment with 10μM BDTP for 10min, the H_2_DCF-DA fluorescence of the BDTP without light group was stronger (39.02%) than that of the control group. And the generation of intracellular ROS in the BDTP with white light irradiation (15mWcm^−2^,30min) was more than other treatment groups. This is because the excited electrons of BDTP can migrate to the lowest triplet excited state, which interacts with ground-state O_2_ (^3^O_2_) to produce ^1^O_2_. Then we investigated the abilities of BDTP to destroy DNA in *V. vulnificus*. As shown in figure 6C, the DNA bands gradually weaken with the increase of BDTP concentration without white light irradiation. Moreover, in the light irradiation (15mWcm^−2^,30min) group, the DNA bands was weakened more obviously. This indicates that BDTP caused DNA damage to *V. vulnificus* and exhibited more significant photodynamic damage. It may be related to the DNA-philic activity of BDTP. This may be due to the oxidative damage of BDTP bound DNA due to the generation of ROS induced by molecular transition. In addition, the morphological changes of the bacteria were investigated using SEM imaging. As shown in Figure. 6D, *V. vulnificus* without BDTP treatment maintained a normal stereotypic morphology with intact cell walls. However, after incubation with BDTP without light irradiation, the bacterial surface appeared obvious damage, and the integrity of some bacterial structures was destroyed. Interestingly, after BDTP combined with light irradiation, more bacteria showed serious surface damage, bacterial cell wall shrinkage and rupture. Therefore, BDTP can not only be used as a bacteria DNA probe under imaging conditions, but also directly destroys *V. vulnificus* through rapid ROS production and sustained damage to bacterial DNA, eventually leading to cell rupture.

**Figure 6.**
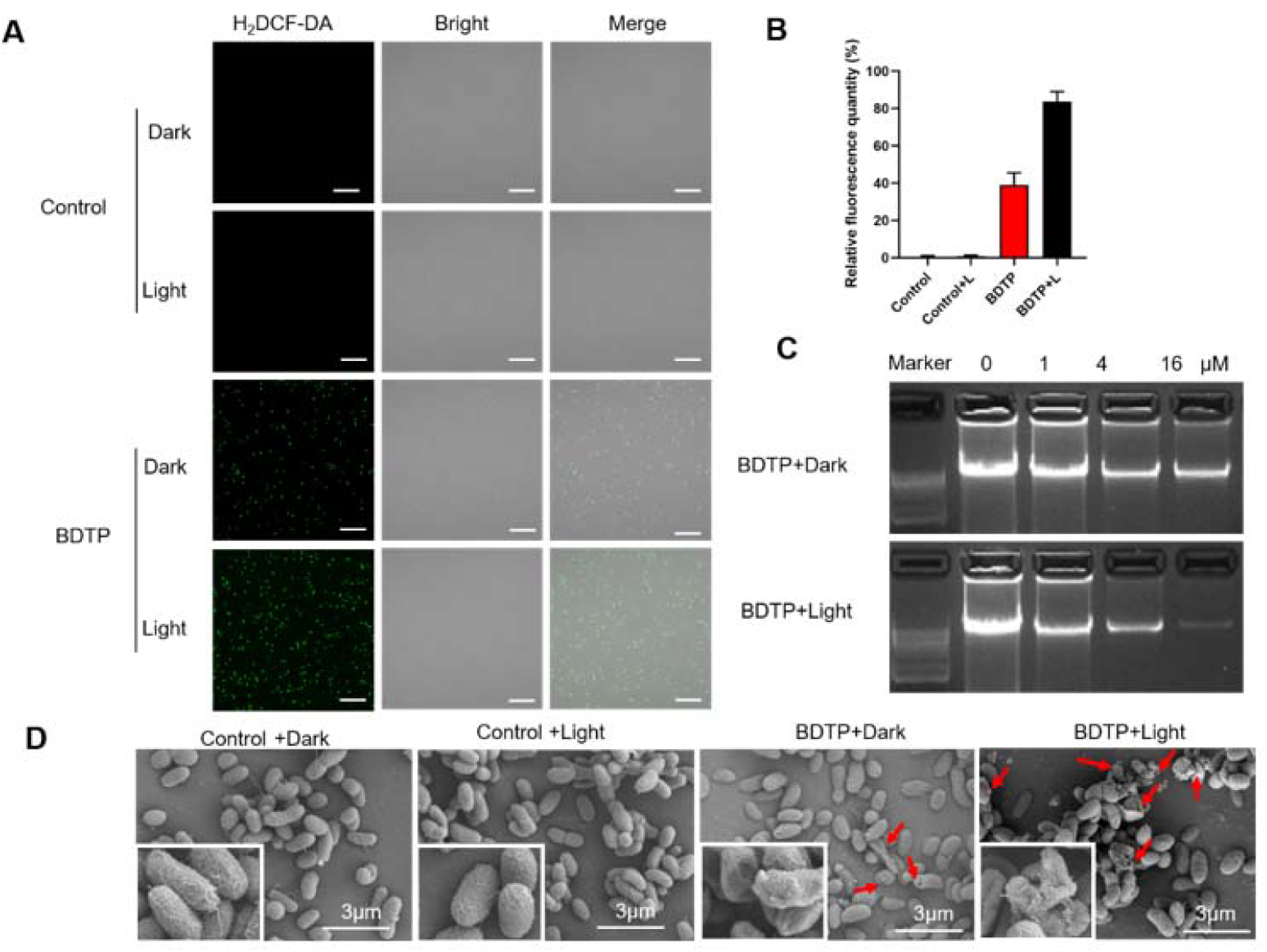
(A) CLSM images of bacteria treated with BDTP. (B)CLSM imaages of *Vibrio vulnificus* incubated with the fluorescent ROS probe H_2_DCF-DA and treated with BDTP (10μM) with or without white light irradiation (15mW cm^−2^, 30min) (n=3 biologically independent samples; mean±SD). (B) Quantitative statistical analysis of fluorescence in figure 5A. (C)Genomic DNA bands of *V. vulnificus* treated with BDTP under different conditions (n=3 biologically independent samples; mean±SD). (D)Scanning electron microscope images of *V. vulnificus* after different treatment with or without light irradiation, and red circle indicate the damage of the bacterial cell wall. Scale bar, 3μm, (n=5 biologically independent samples; mean±SD).

### 2.6 In vivo photodynamic anti-bacterial in wound infection

Based on the above studies, we found that BDTP possesses significant antibacterial and anti-biofilm activities, as well as low cytotoxicity and good blood cell compatibility. Therefore, a BALB/c mouse model of *V. vulnificus* infection was established to evaluate the photodynamic antibacterial effect of BDTP on skin wound infection. Mice full-thickness skin infected wounds were randomly divided into 5 groups: (1) Control group, (2) *V. vulnificus* infection +PBS treatment group, (3) *V. vulnificus* infection + ceftazidime treatment group, (4) *V. vulnificus* infection +BDTP (10μM) with dark treatment group. (5) *V. vulnificus* infection + BDTP (10μM) with light (15mW cm^-2^) treatment group. Took photos of wounds at different time points (1, 3, 5, and 7 days). The schematic diagram depicts the infection as well as the related treatment and observation of the model (Figure 7A). As shown in figure 7B and D, BDTP could inactivate *V. vulnificus* in vivo under dark conditions, and the inactivation efficiency was higher than that of ceftazidime treatment. Moreover, BDTP+L treatment almost completely killed the bacteria in vivo, showing an excellent in vivo sterilization effect. This fully indicates that BDTP has a high antibacterial efficiency in vivo and has efficient photodynamic activity in vivo. In addition, as shown in Figure7C, the wound healing of the BDTP treatment group was better than that of the ceftazidime treatment group, while the wound healing of the BDTP+L treatment group was almost the same as that of the uninfected group. On the third day of treatment, BDTP+L showed a rapid wound healing process, which was better than that of the other treatment groups. In the statistical analysis of the wound area in figure 7E, we can see that the healing rate of the BDTP treatment group reached 39.19% on the 7th day, while the healing rate of the BDTP+L treatment group was as high as 77.72%.

**Figure 7.**
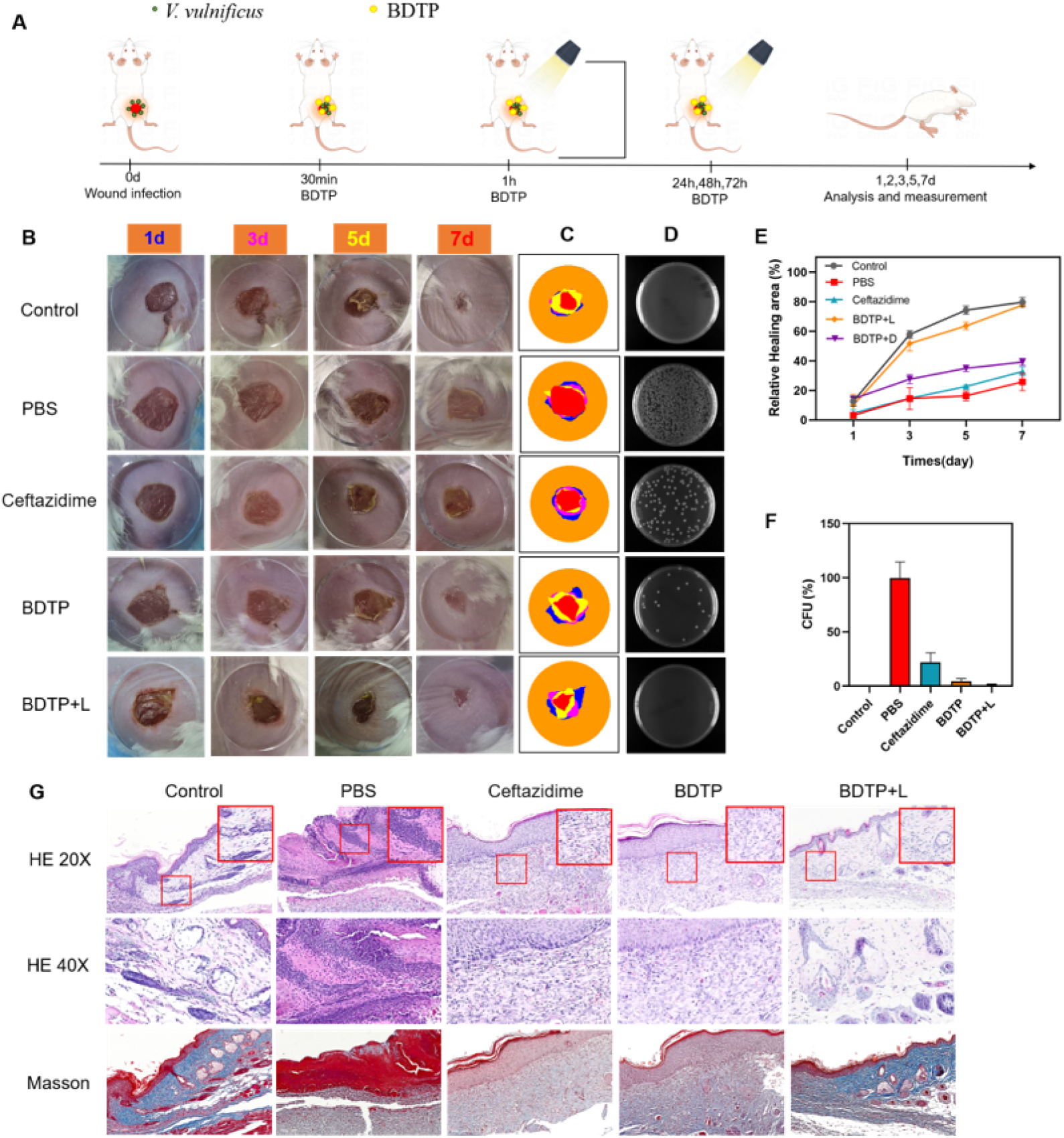
Anti-*V. vulnificus* infection ability of BDTP in vivo. (A) Schematic diagram of *V. vulnificus* wound infection model construction and treatment. (B) Photos of infected wounds of mouse after different treatments at day 1, 3, 5 and 7 (n=5 biologically independent samples; mean± SD). (C) Photographs of bacterial colonies in homogenates of infected sites from mice at day 2 (D) Wound healing rate after infection (n=5 biologically independent samples; mean±SD). (E) Colony culture of tissue homogenates on day 2 of infection (*p<0.05, **p<0.01, ***p<0.001 (n=5 biologically independent samples; mean±SD)). (F) H&E and Giemsa staining of the skin tissue after various treatments (n=5 biologically independent samples; mean±SD).

On day 7 after treatment, the wound tissue of mice was collected for hematoxylin-eosin (he) staining and Masson staining, and the neutrophils generated in the infected tissue were stained blue. As shown in Figure 7F, the number of neutrophils in the PBS-treated group was significantly higher than that in the uninfected control group, and the number of blue neutrophils in the BDTP-only treatment group was significantly less than that in the PBS-treated group and the ceftazidime treatment group, indicating that the latter two groups had more severe wound infection. However, the number of neutrophils in the BDTP+ L-treated group was almost similar to that in the uninfected control group. In addition, neovascularization and fibroblast formation and smooth wound healing were observed in the uninfected control and BDTP+ light group, which may be attributed to bacterial clearance. Wound healing was faster in the BDTP+ L-treated group than in the PBS control group, and at day 7, the latter had much smaller skin lesions and well-shaped hair follicles than the former, which may reflect accelerated dermal growth. Compared with untreated infected wounds, the BDTP+L and BDTP groups significantly promoted hair follicle growth and development, accelerated re-epithelialization of damaged skin, and completely controlled Vibrio vulnificus infection. The photodynamic antibacterial activity of BDTP was quantitatively evaluated by bacterial culture and counting in the tissue around the wound on the second day after treatment. As shown in FIG. 7C, E, the amount of bacteria in the wound of the BDTP treatment group was significantly reduced and bacterial colony formation was less than that of the ceftazidime treatment group, and there was almost no colony formation in the wound tissue of the BDTP+ light group. However, a large number of bacterial colonies were formed in the PBS-treated group. This fully demonstrates that BDTP has excellent antibacterial efficiency in vivo and high photodynamic activity in vivo. In addition, as shown in Figure7C, the wound healing of the BDTP treatment group was better than that of the ceftazidime treatment group, while the wound healing of the BDTP+L treatment group was almost the same as that of the uninfected group.

## 3. Conclusion

We have successfully constructed BDTP, a molecular probe that can localize to bacterial DNA and emit discriminative fluorescence. BDTP showed excellent bacterial indicator function and effective antibacterial ability. In addition, broad-spectrum antimicrobial activity and low toxicity to mammalian cells have been shown. It was used for precise bacterial diagnosis and effective phototherapy in Vibrio vulnificus infected mice. In this study, we developed an effective strategy to construct for the diagnosis and treatment of *V. vulnificus*. Although more and more studies have been conducted on photosensitizers in a wide range of antibacterial effects, and the antibacterial activity of photosensitizers has been significantly improved through different strategies, it is still too early to apply photosensitizers to clinical applications.

## 4. Materials and Methods

### 4.1 Materials

9,10-Anthracenediylbis-(methylene)-dimalonic acid (ABDA), Rose Bengal (RB), Deoxyribonucleic acid sodium salt from calf thymus (DNA (calf thymus)) were purchased from Sigma□Aldrich. Chlorin e6 (Ce6), Dihydrorhodamine-123 (DHR-123) were obtained from Maokang biotechnology (Shanghai, China), Hydroxyphenyl Fluorescein (HPF) was purchased from AAT Bioquest (America). Reactive Oxygen Species Assay Kit (DCFH-DA) was obtained from Solarbio Science & Technology (Beijing, China). Hoechst 33324, Crystal violet dye, CCK8 detection kit and SYTO 9/PI Live/Dead bacterial double stain kit were purchased from Beyotime Biotechnology (Shanghai, China). 2216E nutrient medium was purchased from Haibo Biology (Qingdao, China). The human umbilical vein endothelial cells HUVEC was purchased from Procell Life Technology (Wuhan, China) and cultured in DMEM containing 10% heat-inactivated fetal bovine serum (Gibco, America). *Vibrio Vulnificus* MO6-24/O (*V. vulnificus* MO6-24/O) was obtained from Ho Choi Ph.D. who was the Institute director of the institute of food biotechnology and toxicology nutrition, university of Seoul.

### 4.2 Molecular dynamics simulation

MD simulations were carried out to study the binding and permeation of the BDTP across a hydrated 1, 2-dioleoyl-sn-glycero-3-phosphocholine (DOPC) bilayer.

### 4.3 Bacterial and Cell Culture

*E. coli, S. aureus*, were selected from single clonies, and cultured in lysogeny broth (LB) medium at 37°C and 200 rpm overnight. *V. vulnificus* MO6-24/O was selected from a single clone, and cultured in 2216E medium. By ultraviolet-visible spectra measured at 600 nm in the optical density OD600 to determine the concentration of bacteria. HUVEC cells were cultured in Dulbecco’s modified Eagle’s medium (DMEM; Gibco) with 10% fetal bovine serum (FBS; Gibco) and 1% antibiotic-antimycotic (penicillin and streptomycin) in an incubator at 37 °C with 5% CO^2^.

### 4.4 Antibacterial Activity Measurement

The concentration of bacteria was determined by measuring the OD 600 and adjusted to an OD 600 value of 0.6 by diluting the bacterial suspension with LB or 2216E medium. 1mL bacterial solution was centrifuged at 5000 rpm for 5 min, and washed twice with PBS and resuspended in 1 mL of PBS. 90μL of different concentrations of BDTP solutions were mixed with 10μL of bacterial suspension to achieve final BDTP concentrations of 0, 1, 2, 4, 8, 16, 32, 64, and 128μM, respectively. After an additional 10 min of incubation, the light treated bacteria were irradiated with white light (15mW cm^2^) for 10 min. Then 50μL of the dilution was inoculated on LB AGAR plates or 2216E AGAR plates, and the number of colonies was counted and recorded after 24 h of incubation. All data were carried out in triplicate and presented as mean ± SD.

### 4.5 Antibacterial Activity Assays

In this study, the antibacterial activity of BDTP was examined for *E. coli, V. vulnificus*. MO6-24/O, and *S. aureus*. Single colony of the bacteria was selected and cultured in liquid medium overnight, and the bacterial concentration was adjusted to 1×10^8^CFU mL^-1^. The bacteria were then diluted to 2×10^5^CFU mL^-1^ with normal saline, and BDTP suspensions were prepared with the same buffer. Preparation of different concentrations of BDTP (0,1,2,4,8,16,32,64,128,256 microns) with 1:1 mixed bacteria suspension in 96-well plates. After incubation at 37□ for 10min, the light group was irradiated with white light (15mW/cm^-2^) for 30min,

### 4.6 CCK8 Assays

The HUVEC cell density was adjusted to 1×10^5^/ml, and 100ul of the cell suspension per well was seeded into a 96-well plate and cultured overnight. It was replaced the culture medium containing different concentrations of BDTP (0,1,2,4,8,16,32,64,128μM) according to the method of multiple dilution and incubation 10min. Then the white light (15mW cm^2^) was employed to irradiate for 10 min of the light group. After 24 hours of incubation at 37□, the culture medium was replaced with 100μL fresh medium with10μL CCK8 reaction solution each well and then the absorption at 450nm was measured by microplate reader after continued incubation at 37□ for 2 hours. All data were carried out in triplicate and presented as mean ± SD.

### 4.7 Blood cell compatibility test

1mL blood was obtained from mouse eyeballs, resuspended in 10mL normal saline, and centrifuged at 2000 rpm for 10 min to obtain red blood cells (RBCs). RBCs were washed three times with normal saline, and then 100μL of the red blood cell suspension was mixed with 100μL of different concentrations of BDTP to give a final concentration of 0,1,2,4,16,32,64,128μM. After incubation at 37 °C for 30min, the light group was exposed to white light (15mW cm^−2^) for 30min, followed by centrifugation at 2000 rpm for 10 min. The absorbance of the supernatant at 545nm was recorded using a microplate reader. Triton X-100 (1% v/v) and normal saline were used as positive and negative treatment groups, respectively.

### 4.8 Bacterial and molecular probe fluorescence co-localization

*E. coli, S. aureus, V. vulnificus* MO6-24/O were used for this study. These bacteria were incubated with PBS containing BDTP(4μM) for 10min. And then washed with PBS, followed by co-staining with Hoechst 33324 for 10 min and then the samples were imaged by confocal laser scanning microscopy to investigate the intracellular localization of BDTP.

### 4.9 Fluorescence localization of molecular probes in bacterial and cell co-culture systems

*V. vulnificus* MO6-24/O and the HUVEC cell were used for this study. 80μL of HUVEC cells were seeded in confocal dishes at the concentration of 1×10^5^/ml and cultured overnight,*V. vulnificus*. MO6-24/O was selected and cultured in liquid medium overnight, and the bacterial concentration was adjusted to 2×10^5^CFU mL^-1^. 100μL bacterial suspension was added to the cell culture dish, and then 4μM BDTP was added to the culture dish. After incubation at 37 ° C for 30min, Hoechst 33324 was added and incubated for 10min, and the fluorescence localization of BDTP was observed by laser confocal microscopy.

### 4.10 Scanning electron microscopy Imaging

*V. vulnificus* MO6-24/O concentration diluted to OD600 optical density of 1.0, then the bacteria with 4μM BDTP incubation for 10 min, and then irradiated with 15mW cm^-2^ of white light or dark for 30 min. The cells in the control group were treated with PBS for 30min in the dark. After washing with PBS three times, the bacteria were fixed with 2.5% glutaraldehyde overnight at 4°C. Cells were then dried and subjected to SEM imaging.

### 4.11 Inhibition and Eradication of *V. vulnificus* Biofilms

The inhibitory effect of BDTP on biofilm formation of *V. vulnificus* was detected. *V. vulnificus* suspension (OD600=0.5) and mixtures of different concentrations of BDTP (100μL) were added to 96-well plates and incubated at 37 ° C for 72 hours. The medium was then decanted and washed three times slowly with 200μL of sterile PBS. The bacteria were fixed with 4% paraformaldehyde for 15min and then stained with 0.5% (w/v) crystal violet solution (150μL per well) for 15min. Unbound dye in the Wells was slowly washed three times with sterile PBS. The remaining crystal violet dye of the labeled biofilm was dissolved with 33% acetic acid (200μL/well). The inhibitory effect of different concentrations of BDTP on biofilm formation was quantified by measuring the absorbance value at 590nm. *V. vulnificus* suspension (OD600=0.5) and different concentrations of BDTP mixture were co-cultured for 72 hours, and CLSM was used to observe the biofilm morphology. The staining of SYTO9-labeled biofilms was then observed using CLSM, and the fluorescence was statistically quantified to analyze the inhibitory effect of BDTP on biofilms. The damage effect of BDTP on *V. vulnificus* biofilm was detected. *V. vulnificus* (OD600=0.5) was cultured for 72 hours and then treated with different BDTP. SYTO9/PI was used to co-stain the biofilm, and the staining of biofilm was observed by CLSM.

### 4.12 Laser confocal observation of Intracellular ROS in *vibrio vulnificus*

The ideal photosensitizer has an excellent ability to generate ROS. 2’,7’-dichlorodihydrofluorescein diacetate (DCFH-DA) was oxidized by ROS to produce fluorescent 2,7-dichlorofluorescein (DCF). BDTP was incubated with *V. vulnificus* suspension (OD600=0.5) for 30min, followed by exposure to light or darkness for 30min, and then incubated with DCFH-DA(5μM) for 30min. The PBS treatment group was used as the control group, and the fluorescence intensity of DCF at the maximum emission wavelength of 488 nm was collected by CLSM to evaluate the generation rate of ROS.

### 4.13 Construction of mouse skin injury and infection model

Eight-week-old male BABLC mice were used in the infection model. Sixty mice were equally divided into five groups: (1) the control group without any bacterial infection and additional treatment (n=6); (2) the positive control group was evenly spread on the wound surface with 50 μL bacterial mixture (OD600=0.6) of *V. vulnificus* and then treated with PBS (n=6); (3) the antibiotic-treatment group was evenly spread on the wound surface with 50 μL bacterial mixture (OD600=0.6) of *V. vulnificus* and treated with 16 μM ceftazidime. (4) the BDTP +Dark group was evenly spread on the wound surface with 50 μL of a bacterial mixture (OD 600 = 0.6) of *V. vulnificus* and then treated with BDTP without white-light irradiation (n = 6). (5) the BDTP +Light group was evenly spread on the wound surface with 50 μL of a bacterial mixture (OD 600 = 0.6) of *V. vulnificus* and then treated with BDTP with white-light irradiation (15mW cm^-2^, 30min) (n = 6). All animal procedures were carried out under the guidelines set by the Institutional Animal Care and Use Committee of Shandong Province, and the overall project protocols were approved by the Animal Ethics Committee of the 960th Hospital of the PLA (No. WP20220020).

### 4.14 Histopathology

On day 7, six mice in each group were sacrificed, and the skin tissue was harvested. Skin tissues were fixed overnight in 4% paraformaldehyde, embedded in paraffin, and cut into 5μm thick sections. Hematoxylin-eosin (HE) and Masson trichrome staining were used to evaluate granulation growth, skin maturity, and collagen deposition.

### 4.15 Statistical analysis

Statistical Analysis: All the data were analyzed with GraphPad Prism 8 software and shown as mean ± standard deviation (SD).

---

**Scheme 1.**
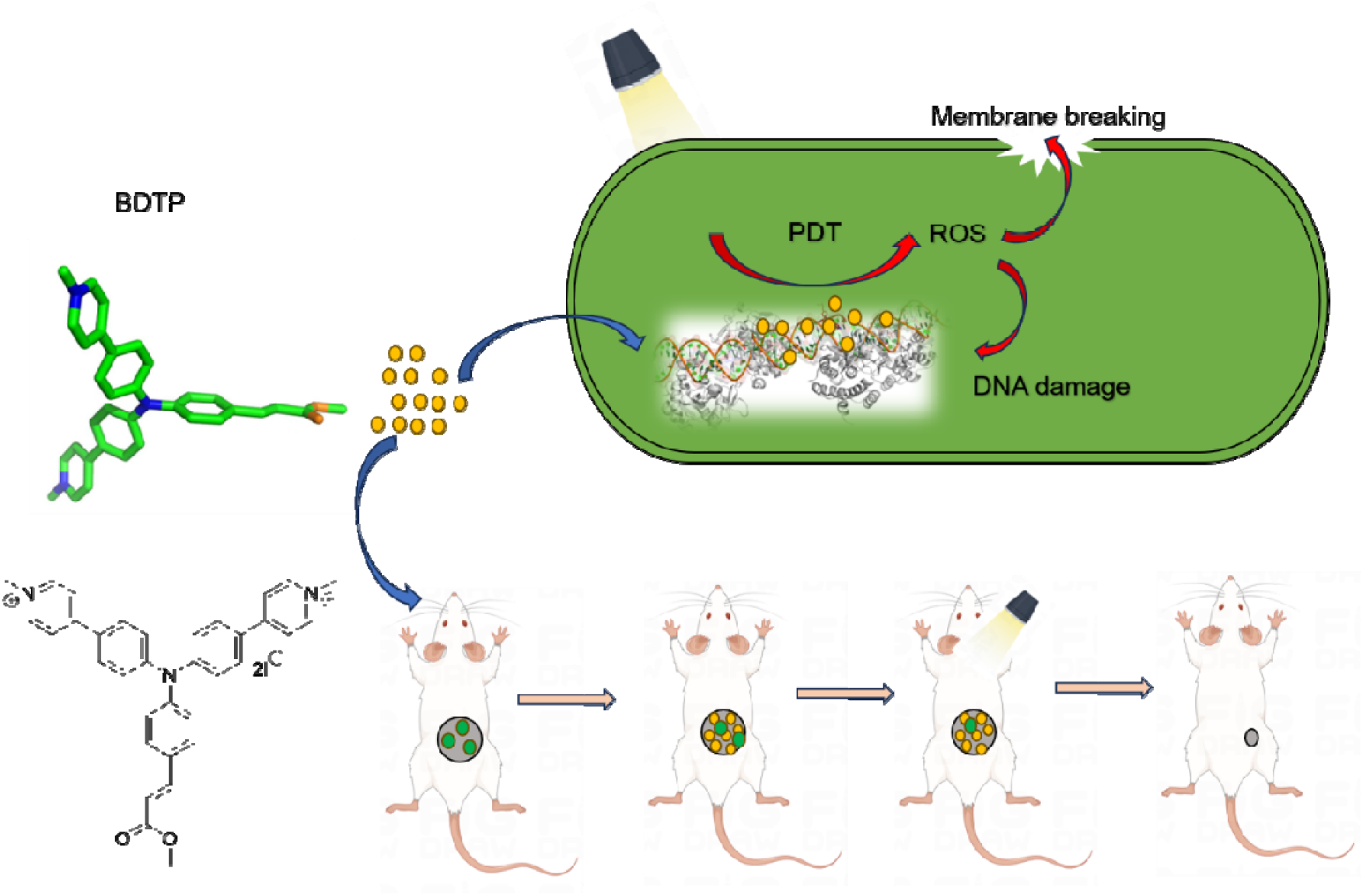
Chemical structure and schematic illustration of BDTP on antibacterial both free bacteria and animal models.

## Notes

### Competing Interest Statement

The authors have declared no competing interest.

